# Pre-task light exposure primes higher-order cognition and preserves mood

**DOI:** 10.64898/2026.06.28.735029

**Authors:** Daniella Mahfoud, Raymond P. Najjar

## Abstract

Light is a fundamental regulator of human physiology and behaviour. Whether prior light exposure shapes subsequent higher-order cognition and mood beyond the period of exposure remains unknown. We tested this in a within-subject, randomised crossover experiment in which 24 healthy young adult males completed a multimodal cognitive battery following 2×15 min of full-spectrum light (FL; median 1,029 melanopic equivalent daylight illuminance [mEDI]) or standard indoor light (SL; median 234 mEDI), with all testing conducted under identical dim illumination. FL improved Digit-Symbol Substitution Test accuracy and promoted digit-directed gaze reallocation, consistent with more efficient associative encoding. On the Balloon Analogue Risk Task, FL reduced reward-seeking behaviour and suppressed backward-referencing gaze transitions linking current and prior-trial reward information. Mood declined following SL but remained stable after FL. Sustained attention, vigilance, and subjective sleepiness were unaffected. Our findings identify pre-task FL exposure as a selective primer of higher-order cognition and mood, independent of alertness.

## Main

Spending time outdoors consistently improves mood, reduces stress, and enhances cognitive performance.^1–3^ These benefits persist after controlling for physical activity, suggesting that sensory features of the outdoor environment, rather than movement alone, contribute to cognitive and affective well-being.^4^ Yet, in modern industrialised societies, people spend around 90% of their waking hours in artificial indoor environments,^5^ making outdoor exposure the exception rather than the norm.

Among the multiple features that distinguish outdoor from indoor environments, light stands out as a particularly potent and quantifiable environmental signal. Daytime outdoor illuminance is generally much higher than indoors, even under shaded or overcast conditions, and can exceed 100,000 lux in direct sunlight, compared with the 100–500 lux typical of indoor environments.^6,7^ Outdoor light is also spectrally broad, rich in short-wavelength energy, and dynamic.^8–10^ Critically, bright indoor light exposure reliably replicates these benefits on mood and alertness in the absence of natural surroundings, ^7,8^ implicating light as a principal environmental signal underlying the psychological and health benefits associated with outdoor exposure.

The retinal gateway for these effects is the intrinsically photosensitive retinal ganglion cells (ipRGCs), which express the photopigment melanopsin, maximally sensitive to ∼480 nm light.^11,12^ Unlike classical photoreceptors, ipRGCs project to brain regions governing arousal, mood, and higher cognition including the locus coeruleus, the lateral habenula, and the prefrontal cortex.^13–15^ Full-spectrum light (FL), engineered to approximate the broad spectral distribution of natural sunlight with high short-wavelength content and high melanopic equivalent daylight illuminance (mEDI), provides a controllable means to engage these pathways and study these effects in the laboratory.

Despite growing interest in the cognitive effects of light, a critical gap remains regarding optimal exposure timing. Previous studies have demonstrated that bright, blue-enriched, and narrowband short-wavelength light can improve alertness, working memory, and prefrontal cortex activation ^16–22^ but these have predominantly delivered light during the task. This design is difficult to implement when tasks must be performed under controlled illumination and poorly reflects real-world behaviour, where light conditions during a task are rarely under one’s control. In contrast, exposure beforehand is readily modifiable: people can step outside before studying, working, or making decisions, then perform these activities under whatever lighting conditions are available. The temporal structure of that exposure may also matter. Intermittent bright light entrains the circadian pacemaker with high efficacy,^23^ and ultrashort light flashes can phase delay the circadian system at least twice as effectively as equiluminous continuous light, yet without inducing corresponding changes in melatonin suppression or alertness. ^24^ Whether this dissociation extends to cognition has never been tested. Whether brief intermittent pre-task exposure to FL can enhance higher-order cognition independently of any light received during task performance remains unknown. Moreover, the field has largely relied on coarse performance measures, overlooking how light reshapes the fine-grained behavioural signatures that reveal *how* cognition operates (*e.g.,* visual scanning strategy and the moment-to-moment gaze patterns underlying task performance).

To address these gaps, we conducted a controlled, within-subject, randomised crossover experiment in which healthy young adults completed a multimodal cognitive battery following brief intermittent (2 x 15 min) exposure to either bright FL (median 1,078 lux, 1,029 mEDI) or standard indoor light (SL, 383 lux, 234 mEDI). Critically, all cognitive testing was performed under identical dim illumination (∼40 lux) irrespective of prior exposure, isolating the lasting effects of pre-task light exposure from the acute effects of light during task performance **(Fig. 1a).** Combining high-frequency eye tracking with conventional behavioural measures, we asked not only whether pre-task FL enhances higher-order cognition, but also how such effects emerge through changes in visual exploration and cognitive strategy.

**Fig. 1:**
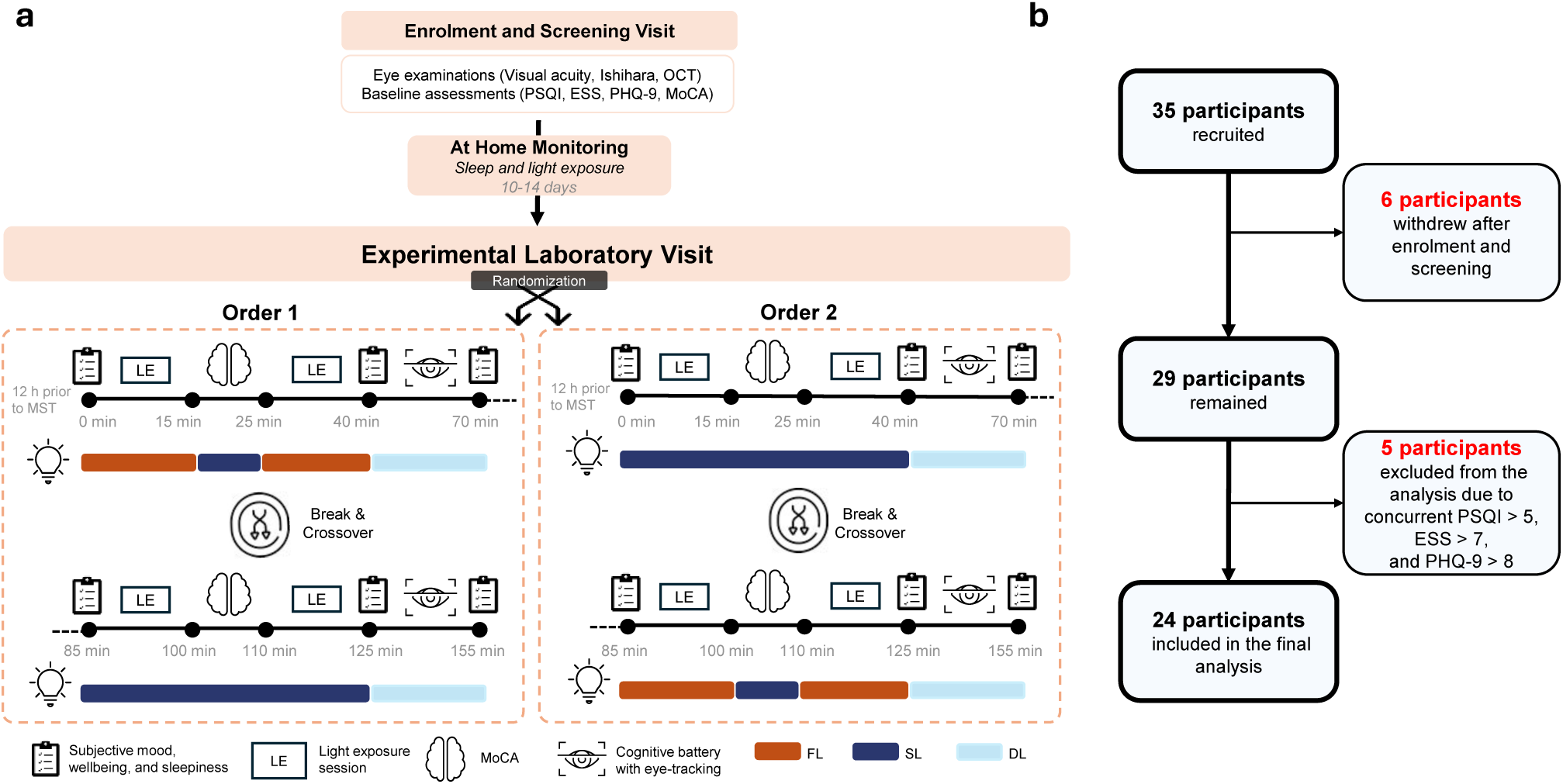
Study design and participant flow. **(a)** Schematic of the experimental protocol. Following enrolment and 10–14 days of at-home actigraphy monitoring, participants completed a single experimental visit in which two counterbalanced light exposure sequences were separated by a break and crossover. In Order 1, full-spectrum light (FL; filled orange bars) was delivered first, followed by standard light (SL; filled dark blue bars); in Order 2, the sequence was reversed. Participants were randomized into orders 1 or 2. Dim LED light (DL; light blue bars) was maintained throughout cognitive testing. LE, light exposure session (15 min). **(b)** Participant flow diagram. Of 35 recruited participants, 6 withdrew after enrolment and 5 were excluded from the analysis for concurrently elevated PSQI > 5, ESS > 7, and PHQ-9 > 8, yielding a final sample of N = 24. *LE, light exposure; FL, full-spectrum light; SL, standard indoor light; DL, dim light; ESS, Epworth Sleepiness Scale; MoCA, Montreal Cognitive Assessment; MST, Mid-sleep time; PHQ-9, Patient Health Questionnaire-9; PSQI, Pittsburgh Sleep Quality Index*.

## Results

### Participant characteristics

Twenty-four healthy male adults (mean age ± SD: 28.2 ± 4.4 years) were included in the analysis (**Fig. 1b**; **Table 1)**. All participants had intact colour vision, normal or corrected-to-normal visual acuity, normal average retinal nerve fibre layer thickness, and low obstructive sleep apnea risk (Table 1). Baseline questionnaires profiles indicated normal cognitive function, higher normal daytime sleepiness (*i.e.,* ESS score range: 6-10), normal sleep quality, and minimal depressive symptomatology (Table 1).

**Table 1.**
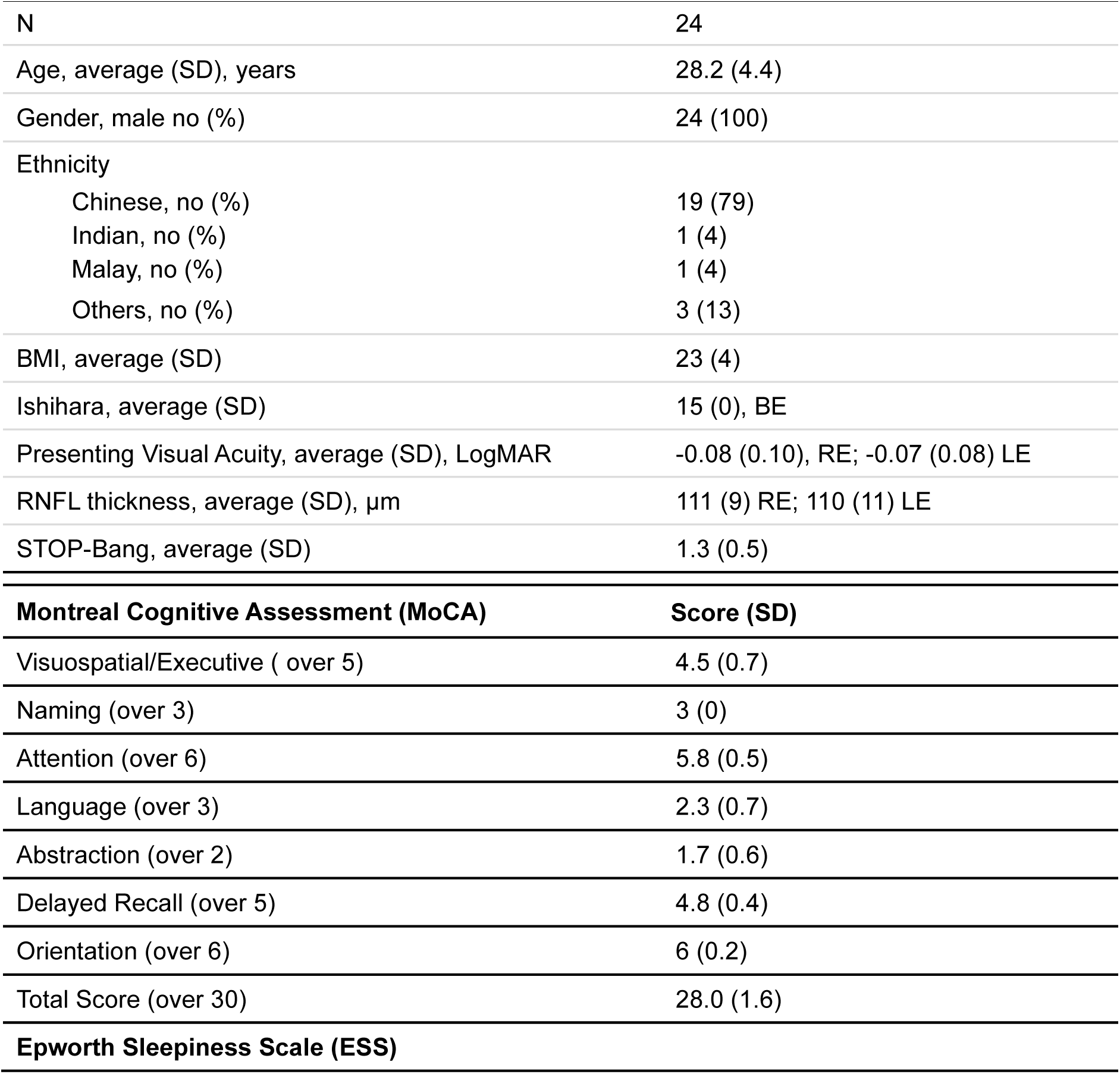
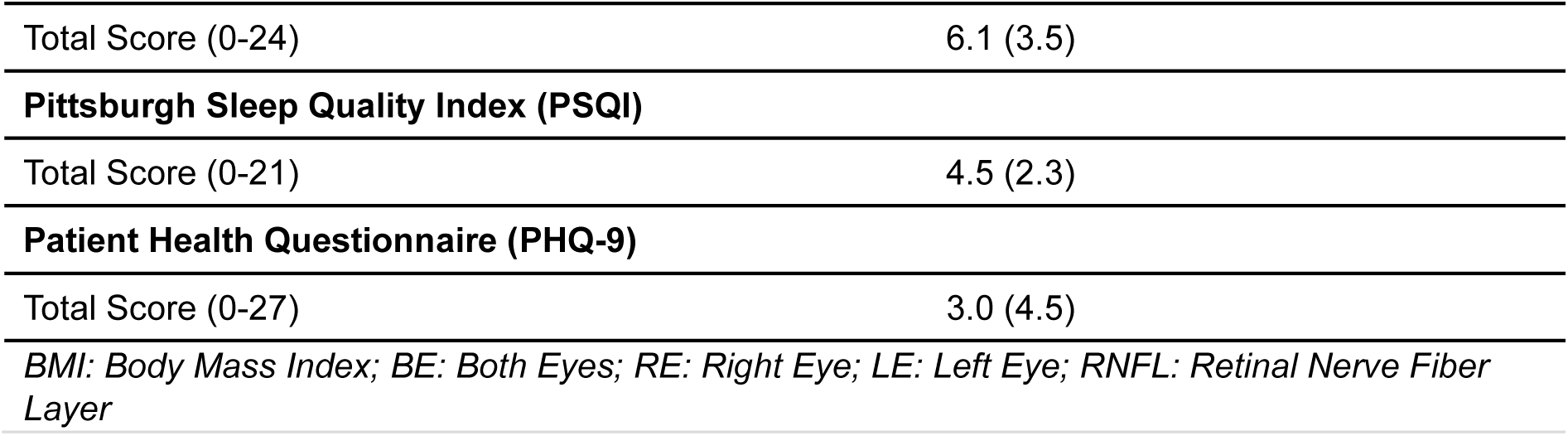
Demographics, clinical, and baseline neurocognitive and behavioural characteristics of participants.

### Pre-visit sleep and light exposure

Actigraphy-derived sleep and light exposure were measured for 10 to 14 days prior to the laboratory visit. Participants exhibited a median mid-sleep time (MST) of 04:47 [04:16–05:34]. The experimental laboratory visit was scheduled to start 12h prior to each participant’s MST to control for potential diurnal impact on outcome measures. Habitual sleep duration was within normal range, with a median total sleep time of 6.5 [6.2–6.9] h and sleep efficiency of 88.5 [86.0–91.2] %. Moderate sleep fragmentation was reflected by a median of 19.7 [12.6–24.4] awakenings per night and 28.1 [18.8–38.7] min of wake after sleep onset (WASO). Participants spent 96.1 [56.6–128.3] min outdoors daily, with a median daytime (between 7:00 and 19:00) illuminance of 204.3 [146.1–298.9] lux and melanopic equivalent daylight illuminance (mEDI) of 169.0 [126.6–286.5] lux. Pre-sleep light exposure (mEDI: 27.6 [16.5–44.9] lux), and light during the sleep period were low (mEDI: 0.9 [0.2–6.1] lux; **Supplementary Table 1**). Real-life sleep and light parameters were not different between counterbalanced order groups (all, *p* > 0.05).

### Pre-task exposure to FL improves Digit-Symbol Substitution Test accuracy and shifts visual strategy toward more efficient digit-directed processing

The Digit-Symbol Substitution Test (DSST) requires participants to check the matching between a target digit–symbol pair and a nine-item reference key. Concurrent eye tracking enabled assessment of gaze-based attentional strategies across the stimulus display.

Accuracy in performing the DSST was improved in 14 out of 24 participants following exposure to FL, compared to SL. Conversely, only 6 participants showed a decline in accuracy and 4 showed unchanged accuracy following FL. Overall, accuracy was higher following FL with a median within-participant change of +1.7% [IQR 0.0 to +3.2%] (Wilcoxon signed-rank: *W =* 52, *p =* 0.046; **Fig. 2a**). The magnitude of improvement was partially constrained by ceiling effects: 9 participants reached 100% accuracy under each condition. Response times did not differ between conditions (SL: 1222 ms [IQR 992–1323] vs. FL: 1217 ms [IQR 1006–1356]; *W =* 143, *p =* 0.86). Accuracy increase was independent of session order (Mann–Whitney U, *p =* 0.09).

**Fig. 2:**
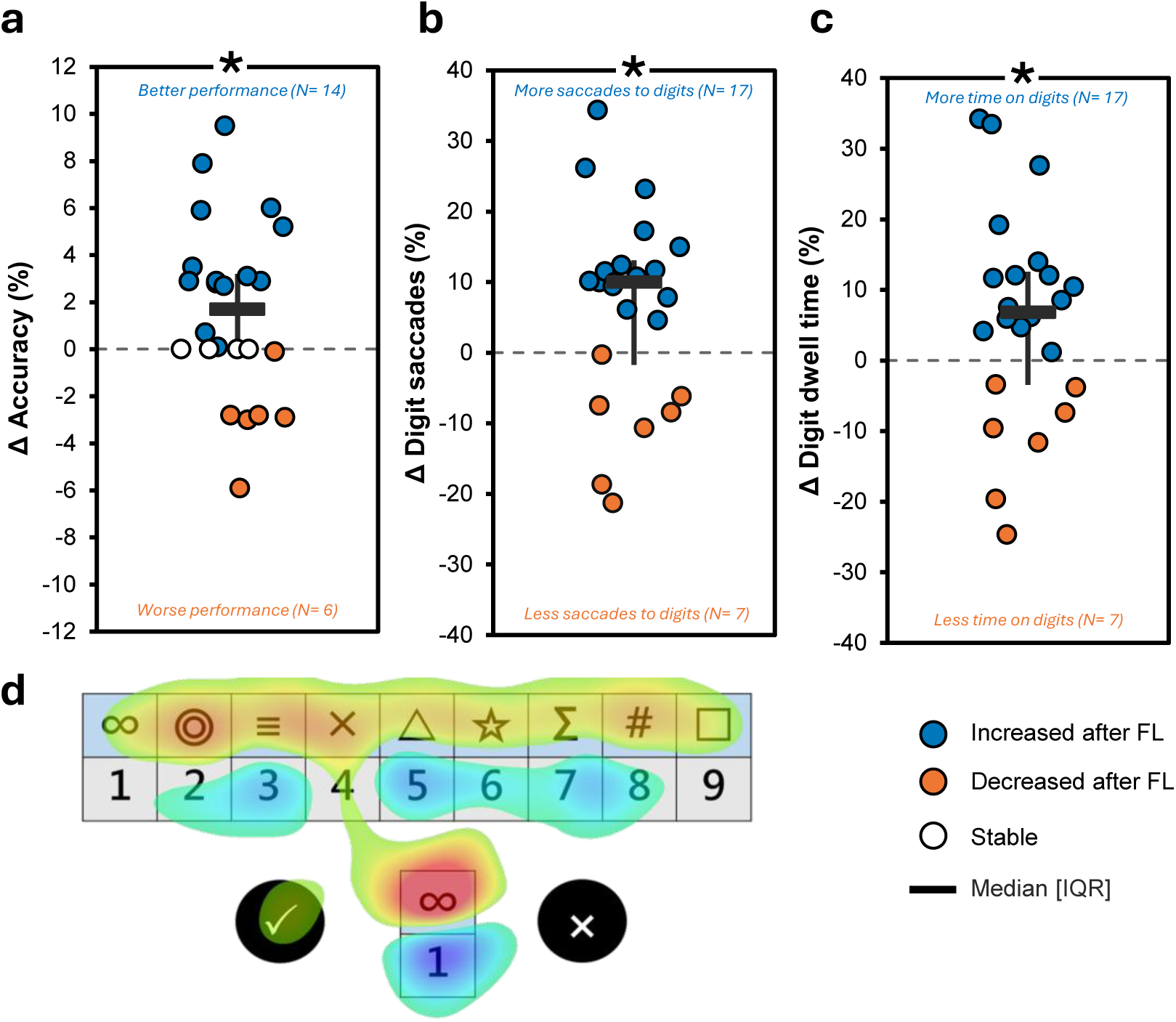
DSST performance and gaze-based visual strategy following exposure to FL or SL. Individual within-participant differences (FL minus SL) in **(a)** accuracy, **(b)** proportion of saccades directed to digit interest areas, and **(c)** digit-directed dwell time. Filled blue circles indicate participants who increased after FL; filled orange circles indicate participants who decreased; filled white circles indicate no change. Horizontal bars indicate the median and vertical lines the interquartile range. Counts of participants showing each direction of change are annotated within panels. **(d)** Representative gaze fixation heatmap during the DSST, illustrating allocation of visual attention across the digit–symbol reference key, response options and stimulus area. The orange areas indicate the regions where there was a reduction in fixations following FL, whereas blue areas indicate the regions where there was an increase in fixations following FL. *\* p < 0.05, Wilcoxon signed-rank test; N=24. DSST, Digit–Symbol Substitution Test; FL, full-spectrum light; IQR, interquartile range*.

Overall fixation behaviour was comparable between conditions: fixation count (SL: 6.5 [IQR 5.6–7.4] vs. FL: 6.6 [5.6–7.2]; *W =* 145, *p =* 0.90), median fixation duration (SL: 162.0 [151.8–169.3] ms vs. FL: 158.0 [149.8–168.5] ms; *W =* 135, *p =* 0.68), saccade count (*W =* 138, *p =* 0.72), saccade amplitude (*W =* 108, *p =* 0.24), and the number of visited interest areas (*W =* 124, *p =* 0.67) did not differ between conditions.

On the other hand, FL produced a systematic shift in the spatial allocation of gaze. Following FL, participants directed a greater proportion of saccades toward digit-related interest areas (SL: 35.0% [IQR 28.3–50.2%] vs. FL: 50.0% [42.1–56.6%]; median within-participant change: +10.0% [IQR −1.7 to +13.1%]; *W =* 67, *p =* 0.02; 17/24 participants; **Fig. 2b**), with a corresponding reduction in symbol-directed saccades. At the subregion level, saccades to the digit grid increased (median change Δ: +6.0%, *p =* 0.046) and to the symbol grid decreased (Δ = −6.8%, *p =* 0.04), while saccades targeting the main digit and symbol items did not differ significantly (both *p* > 0.05).

Digit-directed dwell time increased from 31% [IQR 24–41%] following SL to 41% [36–52%] after FL (Δ = +7.0%; *W =* 76, *p =* 0.03; **Fig. 2c**), and digit-directed fixation count increased similarly (SL: 32% [26–42%] vs. FL: 41% [36–50%]; Δ = +6 pp; *W =* 78, *p =* 0.04; **Fig. 2d**). Subregion analysis revealed that the shift was driven primarily by a reduction in engagement with the main symbol stimulus: dwell time on the main symbol decreased following FL (SL: 27% [24–33%] vs. FL: 25% [20–29%]; *W =* 75, *p =* 0.03), as did fixation count (SL: 28% [24– 33%] vs. FL: 24% [22–29%]; *W =* 81, *p =* 0.049).

### Pre-task exposure to FL reduces risk/reward ratio and suppresses reward-comparison gaze transitions

The Balloon Analogue Risk Task (BART) measures reward-seeking under uncertainty. Participants inflate a virtual balloon across 30 mandatory trials to accumulate earnings, risking losing the trial earnings if the balloon bursts. They may additionally play beyond 30 trials, providing a measure of voluntary reward pursuit. Gaze was recorded continuously to assess task-related information-seeking behaviour.

Prior exposure to FL produced a 62.7% reduction in the risk/reward ratio (adjusted average number of pumps/ total dollars earned across 30 trials) compared to SL (SL: 1.18 [1.11, 1.57]; FL: 0.44 [0.32, 0.58]; *W =* 0, p < 0.001; **Fig. 3a**), with every participant showing a lower ratio following FL. Participants also chose to play fewer total trials after FL exposure (SL: 43.0 [35.8, 51.5]; FL: 36.0 [33.0, 43.3]; *W =* 46, *p =* 0.03; **Fig. 3b**), reflecting reduced voluntary reward-seeking beyond the mandatory 30-trial phase.

**Fig. 3:**
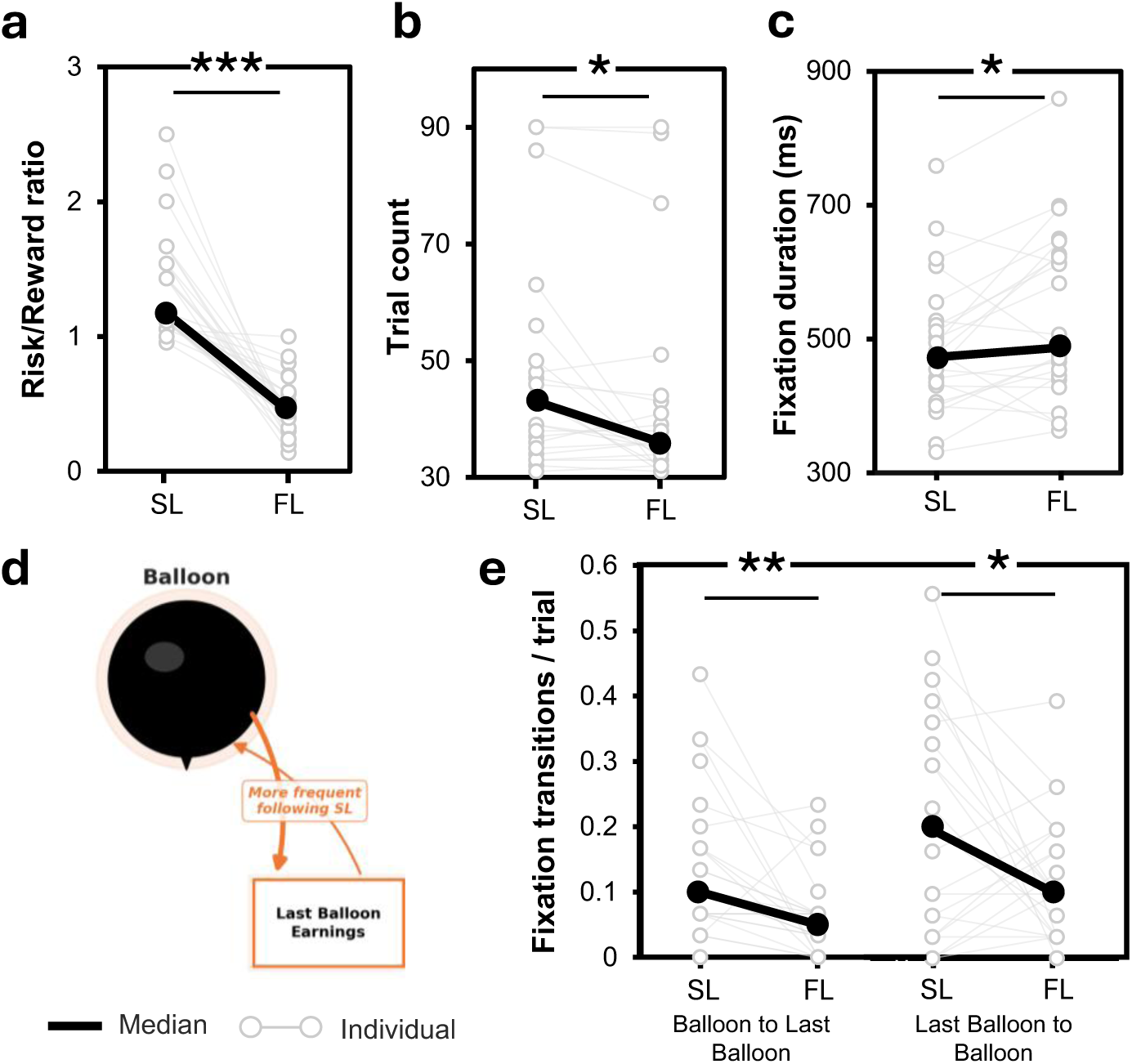
BART risk-taking behaviour and reward-comparison gaze following exposure to FL or SL. Within-participant differences between FL and SL during the Balloon Analogue Risk Task (BART) **(a)** for risk/reward ratio, **(b)** trial count, and **(c)** fixation duration. **(d)** Representative BART display with gaze transitions between the balloon and earnings from last balloon. **(e)** Paired individual data for Balloon Inner → Last Balloon and Last Balloon → Balloon Inner gaze transition rates per trial, respectively. Each paired line represents one participant. Thick black lines connect group medians. *\* p < 0.05; ** p < 0.01; *** p < 0.001, Wilcoxon signed-rank test; N = 24. FL, full-spectrum light; SL, standard indoor light*.

Eye-tracking during the mandatory 30-trial phase revealed longer fixation durations after FL exposure (SL: 473 [424, 522] ms; FL: 487 [443, 623] ms; *W =* 69, *p =* 0.02; **Fig. 3c**). Participants after SL also made more frequent direct gaze transitions between the current balloon and the preceding trial’s earnings display — a reward-comparison circuit quantified as Balloon to Last Balloon transitions (SL: 0.142 [0.072, 0.208] per trial; FL: 0.062 [0.021, 0.094]; *W =* 17, *p =* 0.001) and reciprocal Last Balloon to Balloon transitions (SL: 0.217 [0.089, 0.281]; FL: 0.128 [0.063, 0.172]; *W =* 61, *p =* 0.03; **Fig. 3d-e**), with total dwell in Balloon to Last Balloon transitions likewise reduced (SL: 27.2 [9.8, 36.6] ms; FL: 13.4 [2.8, 19.4] ms; *W =* 48, *p =* 0.01). These gaze-transition patterns are illustrated in **Fig. 3d**. One gaze transition measure (Last Balloon to Balloon) did show a significant order effect (p = 0.003), which should be interpreted with caution.

### Pre-task exposure to FL does not affect sustained and selective attention or temporal perception

Sustained attention (auditory PVT median RT: SL 689 [671, 732] ms; FL 687 [674, 740] ms; *W =* 140, *p =* 0.79, **Fig. 4a**), selective attention (Oddball accuracy: SL 99.4 [98.6, 100.0]%; FL 100.0 [98.6, 100.0]%; *W =* 60, *p =* 0.68, **Fig. 4b**; median RT: SL 730 [695, 775] ms; FL 716 [688, 769] ms; *W =* 130, *p =* 0.57), and time perception (Time estimation task (TET) absolute sum error: SL 15.72 [8.35, 43.07] s; FL 16.35 [10.14, 34.79] s; *W =* 140, *p =* 0.79, **Fig. 4c**) did not differ between conditions.

**Fig. 4:**
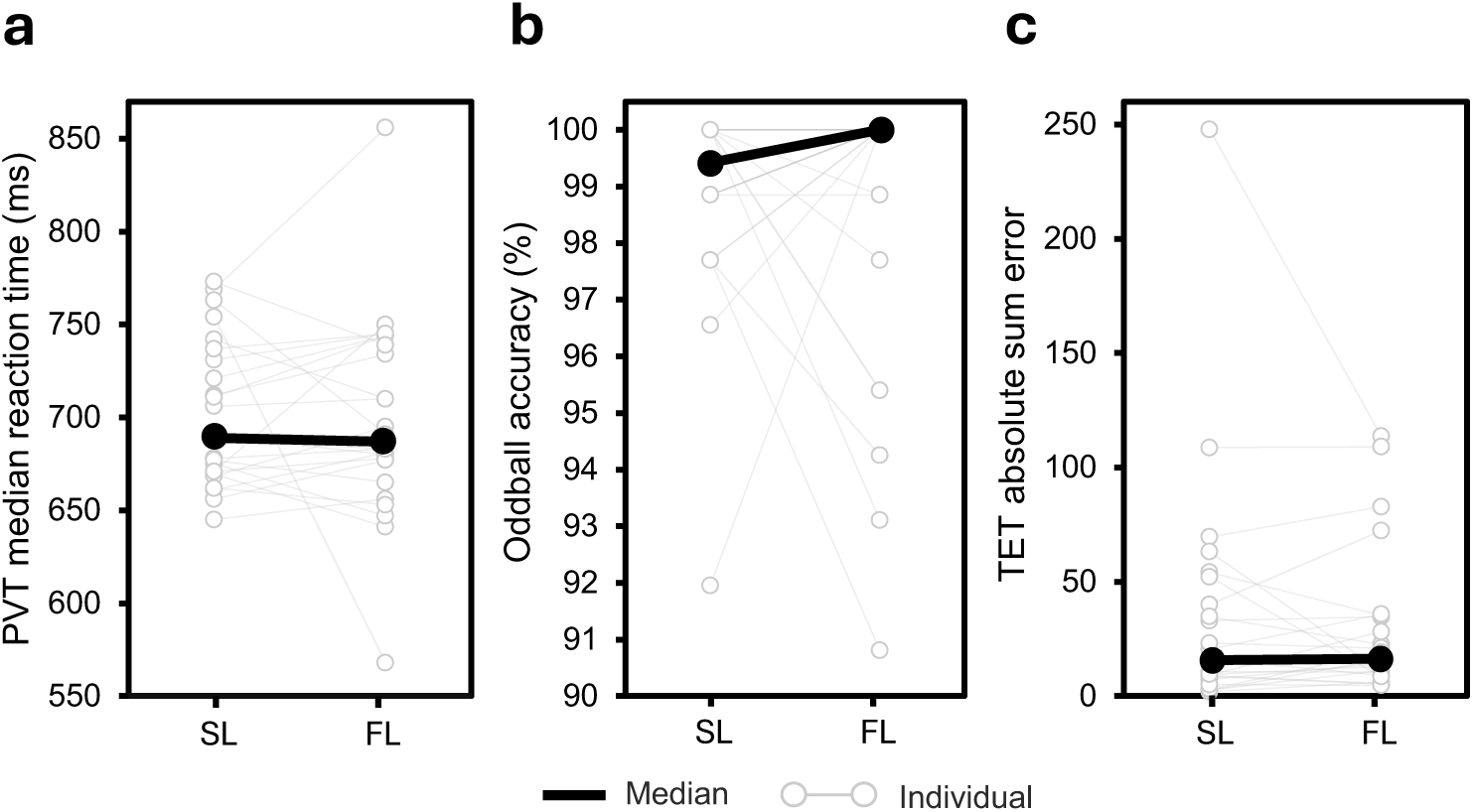
Sustained attention, selective attention and time perception following exposure to FL or SL. Within-participant differences between full-spectrum light (FL) and standard indoor light (SL) for **(a)** auditory Psychomotor Vigilance Task median reaction time, **(b)** auditory oddball accuracy, and **(c)** time estimation task sum absolute error. Each paired line represents one participant and thick black lines show the median. *No significant differences were observed between light conditions, Wilcoxon signed-rank test; N = 24. FL, full-spectrum light; PVT, Psychomotor Vigilance Task; SL, standard indoor light; TET, Time Estimation Task.*

### Prior exposure to FL selectively improves executive but not attentional MoCA domains

The Montreal Cognitive Assessment (MoCA) was administered during the inter-exposure break to assess various cognitive domains. Prior exposure to FL selectively improved the MoCA visuospatial/executive subscale (FL: 4.5 [4.0–5.0] vs. SL: 4.0 [3.0–5.0]; *W =* 4.5, *p =* 0.01). No significant differences were observed for attention (FL: 6.0 [6.0–6.0] vs. SL: 6.0 [6.0–6.0]; *W =* 2.5, *p =* 0.32), delayed recall (FL: 5.0 [4.0–5.0] vs. SL: 4.5 [4.0–5.0]; *W =* 42.0, *p =* 0.28), abstraction (*W =* 2.0, *p =* 0.56), orientation (*W =* 0.0, *p =* 0.32), or language (FL: 2.0 [2.0–3.0] vs. SL: 2.0 [1.0– 2.0]; *W =* 27.5, *p =* 0.06; **Fig. 5**). Total MoCA score did not differ between conditions (FL: 27.5 [26.0–28.2] vs. SL: 27.0 [25.8–28.0]; *W =* 65.5, *p =* 0.223).

**Fig. 5:**
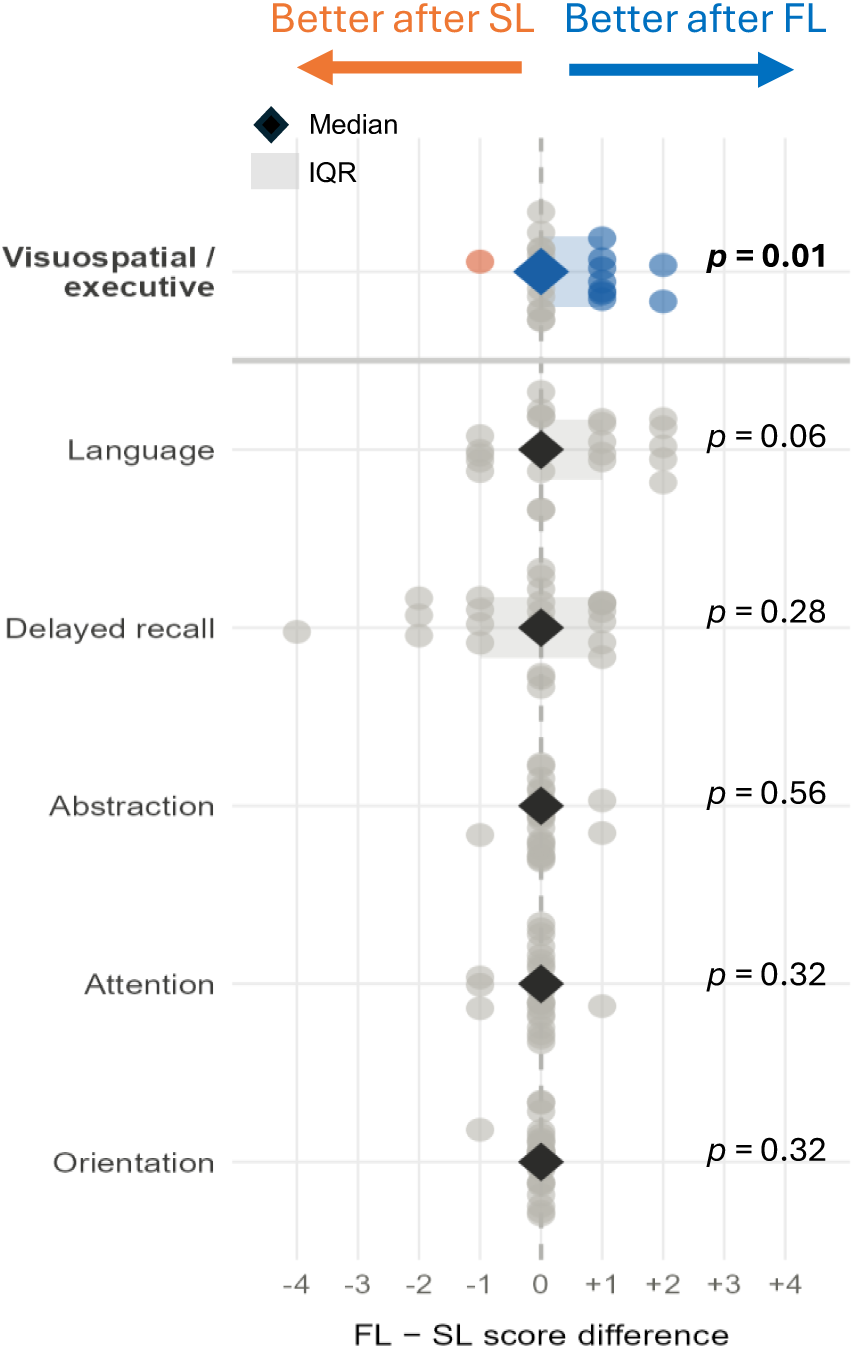
MoCA domains following exposure to FL or SL. Within-participant differences in Montreal Cognitive Assessment (MoCA) domain scores between full-spectrum light (FL) and standard indoor light (SL). Positive values indicate better performance after FL, whereas negative values indicate better performance after SL. Circles represent individual participants, diamonds show the median and shaded bars indicate the interquartile range. Blue and orange points highlight significant individual improvements after FL and SL, respectively, Wilcoxon signed-rank test; N = 24. *FL, full-spectrum light; IQR, interquartile range; MoCA, Montreal Cognitive Assessment; SL, standard indoor light*.

Comparisons against screening baseline (**Supplementary Table 2**) showed that performance following exposure to FL was comparable to participants’ own pre-intervention baseline across most MoCA domains (all *p >* 0.05), whereas SL was associated with significant reductions from baseline in executive function, language, and total score (all, *p <* 0.05). Notably, delayed recall declined significantly from baseline under both conditions (*both, p =* 0.03). The specificity of the FL effect to executive function was confirmed by sensitivity analyses showing no significant influence of light session order (*U =* 74.0, *p =* 0.89), MoCA version assignment (*U =* 52.5, *p =* 0.33), or practice effects (*W =* 27.5, *p =* 1.00) on the FL–SL difference.

### Prior exposure to FL preserves mood while SL drives progressive affective decline

Subjective mood, wellbeing, and sleepiness were assessed at baseline, immediately post-exposure (acute), and 30 minutes post-exposure (sustained) using single-item visual analogue scales and the Stanford Sleepiness Scale (SSS). All scores were normalised to individual baselines. A two-way repeated-measures ANOVA with Condition (SL, FL) and Time (Baseline, Acute, Sustained) as within-subject factors was applied to each outcome.

For mood, significant main effects of Condition (*F*(1,23) = 4.43, *p* = 0.046, η²G = 0.05) and Time (*F*(2,46) = 6.49, *p* = 0.003, η²G = 0.07) were observed, alongside a significant Condition × Time interaction (*F*(2,46) = 5.50, *p* = 0.007, η²G = 0.06). Post-hoc comparisons localised the interaction to the sustained timepoint, where mood was significantly lower following exposure to SL compared to FL (median Δ: SL = −0.50 [IQR: −1.0, 0.0] vs FL = 0.00 [0.0, 0.0]; *W* = 10, *p* = 0.03). Within-condition comparisons confirmed a significant decline in mood following exposure to SL from baseline to the sustained timepoint (*W* = 0, *p* = 0.006), with no significant change at any timepoint following exposure to FL (all *p* > 0.79, **Fig. 6a**).

**Fig. 6:**
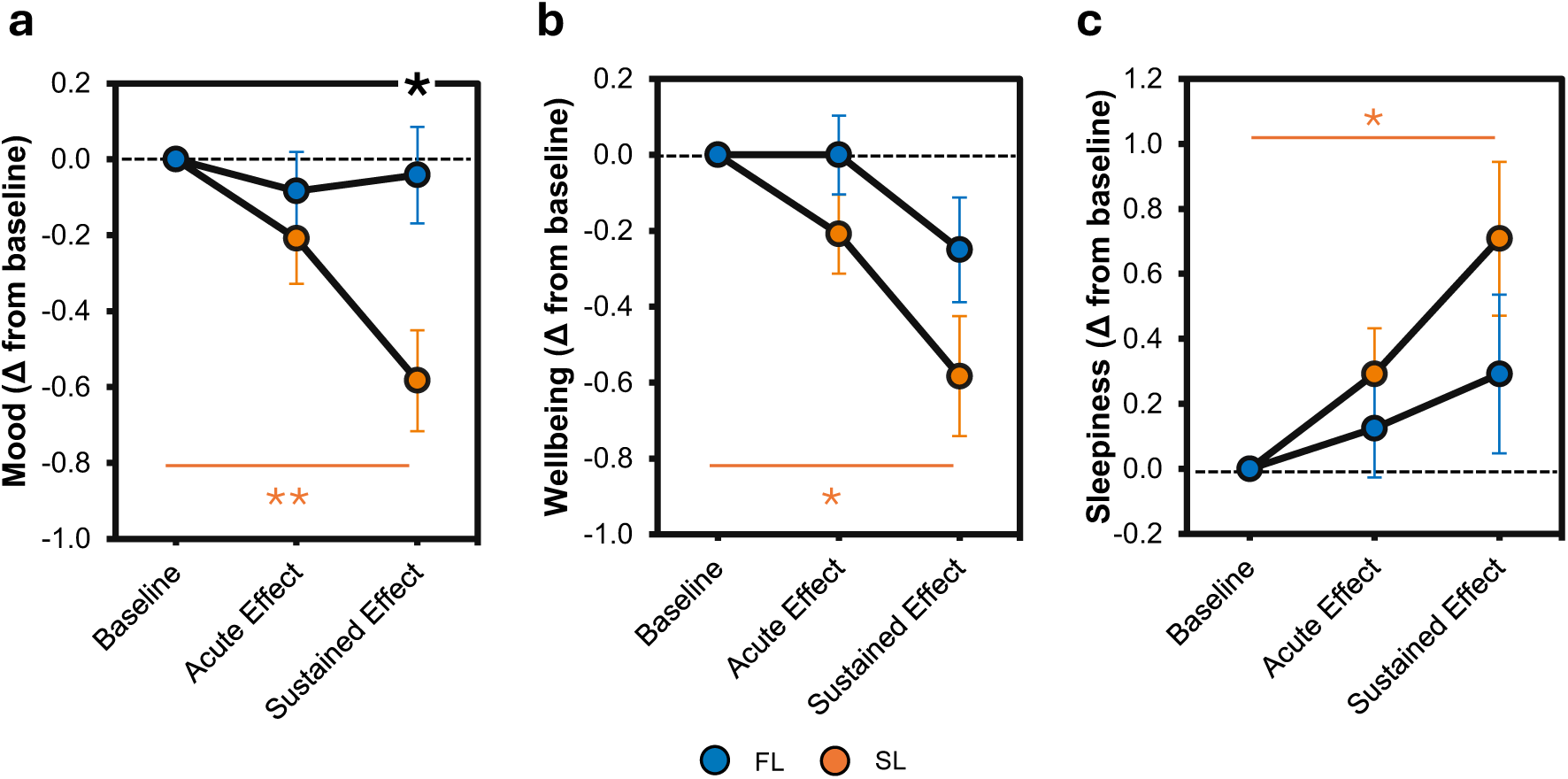
Subjective mood, wellbeing, and sleepiness following exposure to FL or SL. Group means (change from baseline) ± standard error of the mean for **(a)** mood, **(b)** wellbeing (both rated on a 1–7 visual analogue scale; higher scores indicate better state), and **(c)** sleepiness (Stanford Sleepiness Scale, 1–7; higher scores indicate greater sleepiness) at baseline, immediately post-exposure (acute), and 30 min post-exposure (sustained). Filled blue circles represent FL and filled orange circles represent SL. Horizontal orange lines with asterisks denote significant within-SL change from baseline; black asterisk denotes the change between groups at specific timepoint; RM ANOVA and Wilcoxon signed-rank test; *p < 0.05; N = 24. *FL, full-spectrum light; SL, standard indoor light*.

For wellbeing, no significant main effect of Condition (*F*(1,23) = 3.38, *p* = 0.08) or Condition × Time interaction (*F*(2,46) = 1.47, *p* = 0.24) was observed, but a significant main effect of Time was found (*F*(2,46) = 10.23, *p* < 0.001, η²G = 0.11). Post-hoc comparisons indicated a significant decline following exposure to SL from baseline to the sustained timepoint (*W* = 5.5, *p* = 0.02, **Fig. 6b**), with no significant change at any timepoint following exposure to FL (all *p* > 0.05). For sleepiness, no significant main effect of Condition (*F*(1,23) = 1.43, *p* = 0.25) or Condition × Time interaction (*F*(2,46) = 1.30, *p* = 0.28) was observed, but a significant main effect of Time was found (*F*(2,46) = 6.25, *p* = 0.004, η²G = 0.07). Post-hoc comparisons indicated a significant increase in sleepiness following exposure to SL from baseline to the sustained timepoint (*W* = 18, *p* = 0.04, **Fig. 6c**), with no significant change at any timepoint following exposure to FL (all *p* > 0.05). As no Condition × Time interaction was observed for wellbeing or sleepiness, between-condition comparisons were not performed for these outcomes. No order effects were detected for any subjective outcome; change scores did not differ between participants who received SL first versus FL first at either timepoint (all Mann-Whitney U, p>0.25).

Effect sizes across all outcome domains are summarized in **Fig. 7**.

**Fig. 7:**
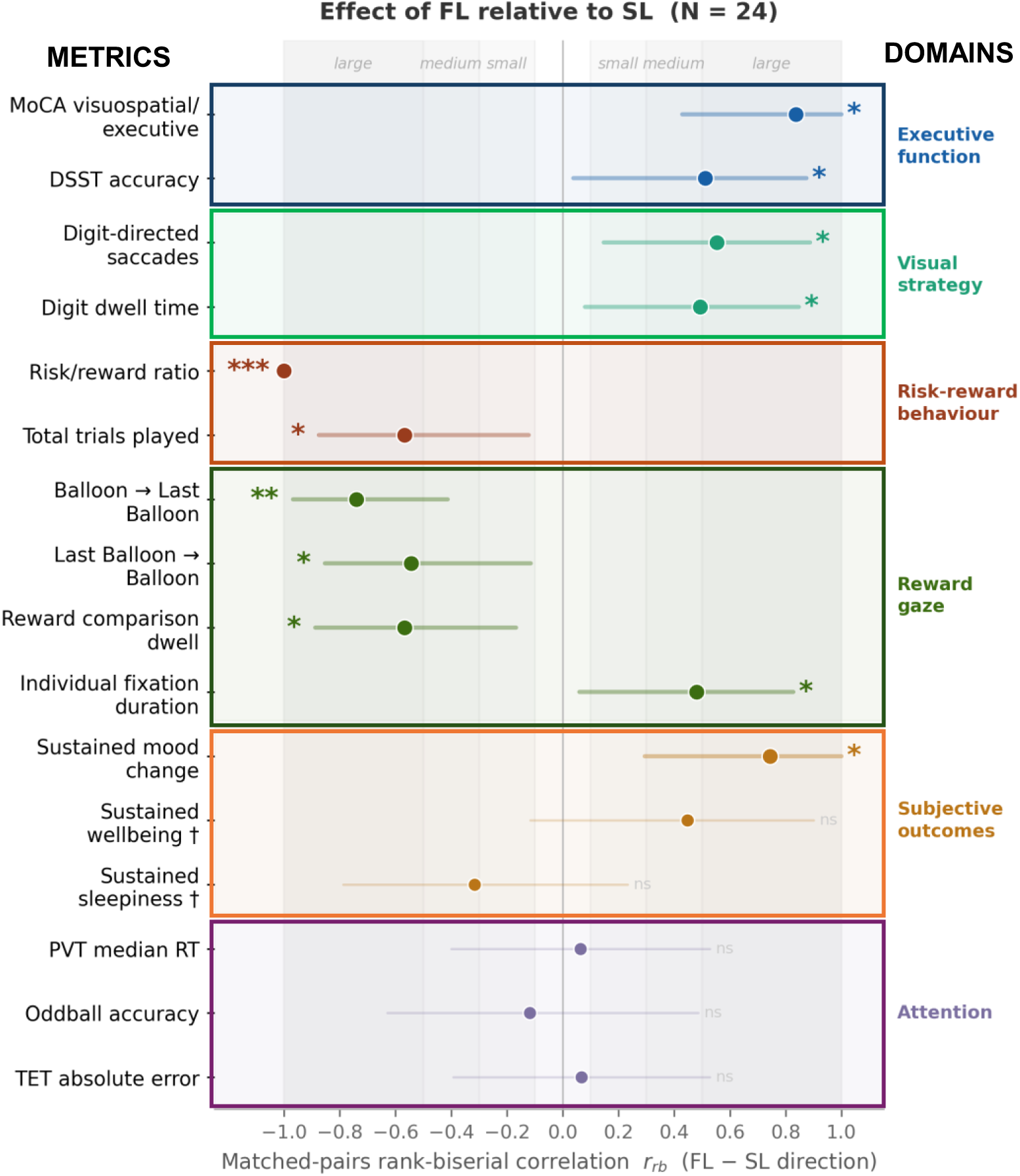
Effect sizes of FL relative to SL across all outcome domains. Each row represents one outcome; the dot indicates the matched-pairs rank-biserial correlation coefficient r_rb_ (computed from the Wilcoxon signed-rank test as ((W^+^ - W^-^)/( W^+^ + W^-^)), and the horizontal bar the 95% confidence interval estimated using paired bootstrap resampling. Positive r_rb_ indicates FL higher than SL; negative r_rb_ indicates FL lower than SL (beneficial direction for risk-reward, gaze-transition, and symbol-engagement outcomes). Rows are organised by cognitive domain and colour-coded. The grey horizontal rule separates outcomes with a significant between-condition difference (above) from those without (below); for sustained wellbeing and alertness (†), SL showed a significant within-session decline whereas FL remained stable, but the between-condition difference did not reach significance (p > 0.12). Shaded background regions denote conventional benchmarks: light grey, small effect (|r_rb_| = 0.10–0.30); medium grey, medium effect (0.30–0.50); dark grey, large effect (> 0.50). N = 24 healthy young adult males. * p < 0.05, ** p < 0.01, *** p < 0.001. *DSST, Digit-Symbol Substitution Test*; *FL, full-spectrum light; MoCA, Montreal Cognitive Assessment; PVT, Psychomotor Vigilance Task; RT, Reaction Time; SL, standard indoor light; TET, Time Estimation Task*.

## Discussion

This study shows that brief, intermittent exposure to bright full-spectrum light can selectively influence subsequent higher-order cognition, reward-guided decision-making, and mood in healthy young adults. By conducting all assessments under identical illumination, we isolated the effects of prior light exposure from the immediate effects of light during task performance.

Our findings extend prior work showing that light can modulate brain activity during cognitive processing.^15^ Neuroimaging studies have demonstrated that the wavelength, duration, and intensity of light exposure can influence brain responses during non-visual cognitive tasks, with particular sensitivity to short-wavelength light.^17,19,22,25–27^ Most relevant to our study, Alkozei et al.^22^ showed that a relatively brief exposure to blue light produced subsequent improvements in working memory performance and temporarily persisting prefrontal activation. We build on this literature in three ways. First, we tested a bright, white, full-spectrum, daylight-like source rather than narrowband blue light. Rather than exposing seated participants to a directed light panel, we illuminated the entire environment using ceiling-mounted full-spectrum panels, with participants encouraged to move during the exposure period. While isolating the contribution of light from other features of the outdoor environment, this setup captures key characteristics of outdoor light exposure—ambient, omnidirectional, and accompanied by physical movement—providing broader stimulation of visual and non-visual photoreceptive pathways than a static, narrowband, directed stimulus. Second, we used an intermittent pre-task exposure design that mirrors how bright light is commonly encountered in daily life through brief episodes of exposure before subsequent activities. Beyond its ecological relevance, this approach was motivated by evidence that intermittent bright light exposure may amplify some non-visual responses compared with continuous illumination.^23,24^ Third, we combined conventional performance measures with high-frequency eye tracking, enabling assessment not only of whether cognitive performance changed, but also of how task behaviour was reorganised after light exposure.

Pre-task exposure to FL enhanced cognitive strategy in addition to improving task outcome. The DSST engages processing speed, visual scanning, working memory, and associative learning, and has been linked to real-world functional abilities.^28,29^ Prior light studies have also reported DSST improvements after blue-enriched or broad-spectrum exposure, supporting its sensitivity to lighting conditions.^30–32^ In the present study, prior exposure to FL improved accuracy without affecting reaction time. However, the effect size was likely constrained by ceiling performance, as 9 participants reached 100% accuracy in each condition. Eye tracking showed increased digit-directed saccades, dwell time, and fixation count, while global oculomotor measures were unchanged. This suggests that participants gazed more selectively at task-relevant information. Because the digits are presented sequentially on the screen, preferential allocation of gaze to the digits may have facilitated digit–symbol matching by reducing unnecessary visual search. Together with the improvement in accuracy, these findings suggest that prior FL exposure modulated task strategy rather than simply increasing general arousal.

Pre-task exposure to FL also influenced reward-guided decision-making. The BART assesses risk-taking by requiring participants to balance increasing reward against increasing probability of loss.^33^ Following exposure to FL, participants showed a lower risk/reward ratio and chose to play fewer voluntary trials beyond the mandatory phase, suggesting reduced reward-seeking or more conservative reward pursuit. This aligns with Lok et al., who reported that broad-spectrum light exposure led to fewer BART pumps compared with dim light, indicating reduced risk-taking.^30^ Eye tracking further showed fewer gaze transitions between the current balloon and previous earnings, indicating reduced visual referencing to recent rewards during ongoing decisions. This suggests that prior exposure to FL changed how participants sampled reward information, reducing reliance on previous reward outcomes during ongoing decisions. In contrast, our findings differ from previous reports suggesting that blue-enriched light increases risk tolerance,^34^ and instead indicate that pre-task FL reduced reward-driven escalation within the BART context.

Our findings suggest that the effects of prior FL exposure were selective and independent of changes in alertness. If these effects simply reflected a generalized increase in alertness, improvements would also be expected in vigilance, sustained attention, reaction time, and subjective sleepiness, none of which differed between conditions. This is consistent with evidence that the alerting effects of daytime light are transient, with improvements in subjective alertness and thalamic and cortical activity declining within minutes after light offset.^27^ Given that cognitive testing in our study was conducted following the final exposure, any acute alertness-related benefits would likely have subsided before testing commenced. This pattern suggests that light can influence higher-order cognitive processes (*e.g.,* associative learning, executive control, visual strategy, and reward-guided decision-making) through mechanisms that persist beyond acute alertness responses. This interpretation is consistent with findings by Lok et al., who reported that 10 hours of daytime broad-spectrum, high-melanopic-efficacy light exposure (100 lux; [mEDI = 80.6 lx]) improved multiple aspects of cognition without significantly altering alertness.^30^

MoCA provides supportive evidence for a selective executive effect. FL improved the visuospatial/executive subscale but not attention, delayed recall, abstraction, orientation, or total score. Given that MoCA is a screening tool rather than an experimental task, this result should be interpreted cautiously. Nevertheless, the domain-specific pattern is consistent with the DSST and BART findings, suggesting selective effects on executive and strategy-dependent behaviour rather than generalized alertness. Beyond cognition, the subjective outcomes suggest that FL may also help stabilise mood during a cognitively demanding session. Sustained mood decreased after standard indoor light but remained stable after FL, while wellbeing and sleepiness did not differ significantly between conditions. Prior work supports the biological plausibility of light effects on mood through melanopsin-sensitive retinal pathways and central projections to regions involved in affective regulation.^13,35^

In the human retina, ipRGCs integrate intrinsic melanopsin phototransduction with rod-and cone-derived signals, exhibit persistent firing after light offset, and constitute the principal retinal input to the suprachiasmatic nucleus as well as other non-image-forming brain centers, ^11,36^ including centers involved in cognitive function and mood such as the prefrontal cortex,^37^ the peri-habenular nucleus.^13^ A plausible interpretation of our findings is that prior exposure to FL, which delivered higher melanopic equivalent daylight illuminance and a broader daylight-like spectrum, elicited greater ipRGC activation than standard indoor light, resulting in more sustained activation of downstream non-image-forming pathways. The absence of changes in alertness despite selective effects on higher-order cognition suggests that different non-image-forming responses may integrate or sustain ipRGC signalling differently. Alternatively, distinct ipRGC subtypes, which project to partially overlapping brain targets,^37–39^ may differentially contribute to alertness and executive functions. Further studies combining behavioural, physiological, and neuroimaging approaches will be required to distinguish between these mechanisms.

The timing of light exposure may be as important as its spectral composition. Most studies investigating the cognitive effects of light deliver illumination during task performance, whereas the present study demonstrates that pre-task exposure alone can produce measurable downstream effects. This distinction has practical implications, as people are often unable to control the lighting conditions under which they work, study, take examinations, or make important decisions, but may be able to influence how they prepare for these activities. Brief exposure to bright, daylight-like light before cognitively demanding tasks may therefore represent a simple, scalable strategy to support higher-order cognition and mood. More broadly, our findings suggest that when light is delivered may be just as important as how it is delivered in shaping cognitive and behavioural responses.

Our study has several strengths, including its within-subject randomised crossover design, identical dim illumination during cognitive testing, pre-visit sleep and light monitoring, personalised scheduling relative to MST, and multimodal cognitive assessment. Most importantly, eye tracking allowed the study to move beyond accuracy and reaction time to identify behavioural signatures of light exposure during task performance. These strengths increase confidence that the observed effects were attributable to differences in pre-task light exposure rather than acute differences in testing conditions. Nevertheless, our findings should be interpreted in the context of several limitations. First, our participants were young and healthy males. Future studies should determine whether these findings generalise to females and older participants who undergo age-related changes in ocular media, non-visual photoreception, and cognitive performance.^40–43^ Second, the 15-min washout between sessions, while preserving feasibility within a single visit, may have been insufficient to fully eliminate carryover in a crossover design. Nevertheless, order effects were non-significant for nearly all outcomes, suggesting that any residual carryover had minimal influence on the observed findings. Third, ceiling effects in DSST accuracy likely attenuated effect size, consistent with evidence that cognitive benefits of light are modest in high-functioning healthy adults.^44^ However, the accompanying eye-tracking findings indicate that changes in visual strategy remained detectable despite this ceiling effect. Fourth, FL and SL differed across multiple dimensions, including illuminance, melanopic EDI, spectral composition, and perceived visual quality. Therefore, isolating which specific characteristic, or combination of characteristics, was responsible for the observed effects remains challenging. Finally, although the behavioural effects were evident, the mechanisms through which FL influenced cognition and mood remain unclear. Combining eye tracking with EEG, autonomic measures, or neuroimaging could help clarify the neural mechanisms through which pre-task light exposure influences higher-order cognition, decision-making, and mood.

In conclusion, brief intermittent pre-task exposure to bright full-spectrum light selectively enhanced higher-order cognition, reward-guided behaviour, and mood stability without affecting alertness. Eye tracking revealed accompanying changes in visual strategy and reward information sampling, suggesting that light modified how participants approached cognitive tasks rather than simply increasing arousal. Together, these findings identify pre-task light exposure as a modifiable environmental intervention that may prepare cognitive and affective systems for subsequent task demands.

## Methods

### Study design

This was a within-subject, randomised, counterbalanced crossover study conducted within a single laboratory visit. The study was approved by the National University of Singapore Institutional Review Board (NUS-IRB-2023-863) and conducted in accordance with the Declaration of Helsinki. All participants provided written informed consent prior to participation.

Each participant completed two light exposure sessions in sequence during a single visit, separated by a 15-min washout period to minimize carryover effects while maintaining feasibility of completing both conditions in one day. The order of light exposure conditions (FL *vs.* SL) was randomized across participants using a computer-generated randomization sequence. Neither participants nor researchers were blinded to condition assignment due to the perceivable difference in light conditions. All cognitive assessments were administered in a standardised manner by a trained examiner.

### Participants

Participants were recruited from the general public. Inclusion criteria required healthy male adults aged 21–35 years with presenting visual acuity (VA) better than 6/9, normal colour vision (Ishihara plates), no history of diagnosed ocular disease (excluding dry eye or myopia ≤5.00 dioptres), no systemic condition or medication affecting pupillary responses or alertness, no clinically diagnosed neurological, psychiatric, or sleep disorders, no night-shift work within the preceding three months, and no recent transmeridian travel (≤ 1 month).

A total of 35 male adults were enrolled. Six participants withdrew before their experimental visit. Five participants were excluded post-randomization based on pre-specified criteria indicating elevated scores across all three of the following measures: Pittsburgh Sleep Quality Index (PSQI) > 5, indicating poor sleep quality; Epworth Sleepiness Scale (ESS) > 7, indicating high level of daytime sleepiness; and Patient Health Questionnaire-9 (PHQ-9) > 8, indicating elevated/moderate depression. The final analysis therefore included N = 24 participants. A CONSORT-style participant flowchart is provided in **Fig. 1b**.

Sample size calculation was informed by the effect of broad-spectrum LED exposure on DSST performance reported by Lok et al.^30^ (d = 1.17, within-subject crossover design). Power analysis based on this effect size indicated a minimum sample of N = 8 (α = 0.05, power = 0.80, two-tailed paired comparison). Given that their exposure contrast (broad-spectrum vs. dim light, 10 lx) and prolonged duration (10 h continuous) likely overestimate the effect size expected under our more conservative FL versus SL comparison with intermittent 2 × 15-minute exposures, we targeted a sample approximately four times larger. After withdrawals, 29 participants remained, with 24 retained following application of pre-specified exclusion criteria (concurrently elevated PHQ-9, ESS, and PSQI scores).

### Pre-visit assessments

An enrolment and screening visit was conducted around two weeks prior to the experimental laboratory visit. During this visit participants provided written informed consent, underwent eye assessments (visual acuity, Ishihara, and optical coherence tomography) and completed a baseline characterisation battery. Risk of obstructive sleep apnea was assessed using the STOP-Bang questionnaire. Participants also completed four validated self-report instruments: the PSQI,^45^ which measures habitual sleep quality and disturbance over the preceding month; the ESS,^46^ which quantifies propensity to doze across daily situations; the PHQ-9,^47^ a validated screen for depressive symptom severity; and the Montreal Cognitive Assessment (MoCA, Singapore English Version 7.1),^48^ a brief standardised tool sensitive to mild cognitive impairment across multiple domains. Except for MoCA, the lower the score, the better the outcome.

At the end of the enrolment visit, participants were fitted with a wrist-worn actigraphy and light monitoring device (ActLumus, Condor Instruments, Brazil) and provided with a sleep diary. Participants wore the ActLumus and filled the diary continuously for 10–14 days.

Actigraphy data served two purposes: to characterise habitual sleep quality and ambient light exposure in the period preceding the laboratory visit, and to determine each participant’s mid-sleep time. The main experimental session was then individually scheduled to begin 12 hours prior to mid-sleep to minimise inter-individual variability in circadian phase at testing.

Actigraphy-derived sleep metrics included: total sleep time (TST), time in bed (TIB), sleep efficiency (SE%), wake after sleep onset (WASO), number of awakenings, and sleep onset latency (SOL). Habitual light exposure was characterised by outdoor time (minutes), average illuminance (lux), and average melanopic equivalent daylight illuminance (mEDI) at different time windows (daytime, pre-sleep (3 hours prior to sleep), and sleep).

### Experimental laboratory visit protocol

All testing sessions were conducted in the afternoon, with timing individualised per participant and set at 12 hours prior to each participant’s mid-sleep time (MST), to minimise potential confounds from circadian phase. Participants were instructed to maintain habitual caffeine consumption on the day of testing. The visit consisted of two counterbalanced sessions. Each session began with a baseline assessment of subjective mood and wellbeing using single-item visual analogue scales (1–7 scale), and sleepiness using the Stanford Sleepiness Scale (SSS).^49^ Participants then underwent 30 minutes of light exposure (2 × 15 min blocks separated by 10 minutes when the MoCA test was administered). During light exposure, participants were asked to walk around a white empty room with the possibility to sit on the chairs in the middle **(Supplementary Fig. 1a-b).** They were permitted to blink naturally but were asked to avoid using their phones, closing their eyes for extended periods, eating, sleeping, or doing any cognitively stimulating activity. Subjective scales were repeated immediately after the second exposure (acute effect) and 30 min later at the end of the session (sustained effect). Following the acute assessment, participants completed a cognitive battery with concurrent eye tracking: DSST, aPVT, Auditory Oddball, TET, and BART under dim light **(Supplementary Fig. 1c-d).** The same procedure was then repeated for the second session (crossover condition). A 15-min break separated the two sessions where participants were allowed to drink water, use the washroom, and rest.

### Light characteristics

Light exposure was delivered using custom-built LED ceiling lights with adjustable intensity and spectral composition (ColorDyne, UK). Two lighting conditions were delivered via ceiling lights. The full-spectrum light source (FL) delivered a median photopic illuminance of 1078 lux [IQR 940-1132] with a correlated colour temperature of 6,205 K, a median melanopic equivalent daylight illuminance (mEDI) of 1,029 lux. The standard light condition (SL) delivered a median photopic illuminance of 383 lux [270-492], a CCT of 3,955 K and a median mEDI of 234 lux (Supplementary Fig. 2).

Light measurements were conducted using a calibrated spectroradiometer (Jeti Specbos 1211, Jeti Technische Instrumente GmbH, Germany) at 150 cm height, facing upwards, across 13 room positions. Melanopic equivalent daylight illuminance (mEDI) was calculated using the CIE S 026 α-opic Toolbox – v1.049a – 2020/11/16.^50^ Spectral irradiance measurements were taken at 13 spatial positions spanning the full depth and width of the room **(Supplementary Fig. 2).**

### Cognitive assessments

#### Montreal Cognitive Assessment (MoCA)

The MoCA is a brief, validated, screening instrument that assesses multiple cognitive domains including attention, executive function, memory, language, visuospatial abilities, and orientation.^48^ Total scores range from 0 to 30, with higher scores indicating better cognitive function. The MoCA was administered and graded by a trained examiner (Certification ID: SGMAHDA710679697-02) according to standard instructions. During the enrolment visit, MoCA Singapore English (version 7.1) was performed using pen and paper. During the experimental visit, alternate validated versions (version 8.2 and 8.3) were used through the MoCA duo app across sessions (randomly assigned version) to minimise practice and order effects.^51,52^ MoCA was done between the 2 light blocks in each session (Fig. 1).

### Post-exposure assessments

A comprehensive cognitive battery was administered after the end of each light exposure session with concurrent eye tracking (Fig. 1). The performance setup consisted of a chair and table with the eye tracker, a screen, a mouse, and a joystick. Participants consistently fixated on the chin rest and had headphones on while performing the tasks. They were exposed to light of LED spectrum at 40 lux at eye-level. The battery included the following assessments, administered in fixed order:

#### Auditory Psychomotor vigilance task (PVT)

PVT is a reaction time task that measures sustained attention and vigilance.^53^ A modified auditory version of the PVT was used in this study. Participants were instructed to fixate on a black circle on the screen and press a button as quickly as possible when they hear a beep through their headphones. The beep appeared at random intervals ranging from 2 to 6 seconds over a 5-minute testing period. The primary outcome measure was median reaction time (RT), with slower reaction times indicating reduced vigilance.

#### Digit-symbol substitution Test (DSST)

The DSST is a paper-and-pencil test of processing speed, working memory, and associative learning.^28^ A modified and computerized version of the DSST was administered in our study. In this version, participants were presented with a reference grid showing nine digit-symbol pairs at the top of the screen, followed by a main digit-symbol pair at the bottom of the screen that changed in each trial. Participants’ task was to identify if the main pair matches the pairs in the reference grid. If so, participants were instructed to press the left arrow key corresponding to “Correct”, otherwise they should press the right arrow key corresponding to “Incorrect”. If a participant took more than 5 seconds to press the arrow key, the trial was considered a lapse. Two versions of the DSST were used randomly and interchangeably to avoid learning and practice effect. The first two trials of the first session were excluded and considered as training sessions. Performance was quantified as the percentage of correct responses (accuracy) and reaction time (speed). High-frequency eye tracking (described below) was conducted during DSST performance to assess visual strategies and gaze behaviour.

#### Auditory oddball task

This task assessed selective attention and inhibitory control.^54^ Participants listened to a series of tones (80% standard tones at 780 Hz, 20% deviant tones at 1000 Hz) through headphones and were instructed to press a button only in response to deviant tones while withholding responses to standard tones. The task included 90 trials separated by 2 seconds. Performance was quantified as accuracy (percentage of correct responses to deviant tones).

#### Time estimation task (TET)

Temporal perception was assessed using a time estimation task administered to monitor sleepiness objectively.^55,56^ On each trial, participants were asked to estimate 2 minutes since the start of the experiment. These 2 minutes were split into checkpoints. Participants were presented with a screen displaying a target duration (10, 20, 30, 60, or 120 seconds) alongside a fixation box and were instructed to press the spacebar when they judged that the specified duration had elapsed since the start of the experiment. Participants were permitted to count. Performance was quantified as absolute sum error of all the durations.

#### Balloon analogue risk task (BART)

The BART is a computerized measure of risk-taking behaviour and decision-making under uncertainty.^33,57^ Participants were presented with a balloon on the screen and could earn money by inflating it with repeated button presses. Each press earned a $0.05 reward, but the balloon could explode at any point, resulting in loss of all earnings for that trial. Participants could alternatively choose to cash out at any time to bank their earnings. The task included 30 main trials with varying explosion probabilities. In this variation of BART, participants were permitted to continue playing beyond the initial 30 trials and stop at their discretion, a modification designed to isolate reward-seeking behaviour in the absence of negative consequences. Participants were given vouchers corresponding to their gains. Primary behavioural outcomes were total trial count, indexing overall reward-seeking propensity, and the risk/reward ratio, defined as the mean number of pumps per dollar earned across collected trials, where higher values indicate greater risk exposure per unit of reward obtained. Concurrent eye tracking was recorded throughout all trials.

### Eye tracking

High-frequency eye tracking was conducted during DSST and BART performance using a remote eye tracker (EyeLink 1000 Plus, SR Research, Canada) sampling at 1000 Hz. The eye tracker was positioned facing the participants. Participants were instructed to maintain natural head position during the task, and a chin rest was used to minimize head movements while allowing comfortable writing posture. Prior to each task, a 9-point calibration procedure was conducted to ensure accurate gaze position estimation. Although binocular eye tracking was recorded, analyses were conducted using data from one eye only. The study eye was selected randomly when both eyes passed calibration and validation; otherwise, the eye that successfully passed calibration and validation was selected for analysis.

For DSST, interest areas (IAs) were defined at four levels: (1) the reference digit grid; (2) the reference symbol grid; (3) the main digit stimulus; and (4) the main symbol stimulus.

For each participant and condition, we calculated: total fixation count per IA, total dwell time per IA, median fixation duration per IA, and number of transitions between IAs (saccade count directed to each IA). For BART, IAs were defined over the current balloon and the preceding trial’s earnings display (Last Balloon) as well as total earnings. Outcomes included fixation duration and transition rates in Balloon–Last Balloon gaze circuits.

### Statistical analyses

All analyses were performed in R (version 4.5.0). Data were assessed for normality using the Shapiro–Wilk test (threshold p < 0.05). Given that most outcomes violated normality, paired comparisons between light conditions were conducted using Wilcoxon signed-rank tests. Effect sizes are reported as matched-pairs rank-biserial correlations, calculated as r_rb_ = (W^+^ - W^-^)/( W^+^ + W^-^), where W^+^ and W^-^ are the sums of positive and negative signed ranks, respectively. Effect sizes were signed in the FL − SL direction, such that positive values indicate higher values under FL than SL and negative values indicate lower values under FL than SL. Ninety-five percent confidence intervals for r_rb_ were estimated using paired bootstrap resampling. Effect magnitudes were interpreted using conventional thresholds: |r_rb_| ≥ 0.10 small, ≥ 0.30 medium, ≥ 0.50 large).

For subjective outcomes (mood, wellbeing, sleepiness), a two-way repeated-measures ANOVA (Condition [FL, SL] × Time [Baseline, Acute, Sustained]) was applied, with generalised eta-squared (η²G) as the effect size. Post-hoc pairwise comparisons used Holm-Sidak-corrected Wilcoxon signed-rank tests. Order effects (FL-first vs SL-first) were tested using Mann–Whitney U tests on within-participant FL–SL difference scores. Session order did not moderate any subjective or primary cognitive outcome. Statistical significance threshold: α = 0.05.

## Author contributions

Concept and design: DM and RPN. Data acquisition: DM. Analysis and interpretation: DM and RPN. Manuscript preparation: DM and RPN.

## Acknowledgments

We would like to thank the research participants who took part in this study as well as T Vicknaswari and Jiaul Baksh who supported the recruitment and screening visit. This research was funded by the ASPIRE-NUS startup grant (NUHSRO/2022/038/Startup/08) to RPN.

## Data availability

The de-identified datasets used in this study can be made available from the authors upon reasonable request. The full study protocol is available from the corresponding author on reasonable request.

## Code availability

The code used in this study can be made available from the corresponding author upon reasonable request.

## Ethical approval and consent for publication

Consent was obtained directly from participants. The study was approved by the National University of Singapore Institutional Review Board (NUS IRB), NUS-IRB-2023-863.

