## Supplementary Material for "Pre-task light exposure primes higher-order cognition and preserves mood"

### This supplementary material document includes:

**Supplementary Figures:**

Supplementary Fig. 1: Experimental setup and light source characterisation

Supplementary Fig. 2: Spatial distribution of melanopic equivalent daylight illuminance (mEDI) across the light exposure room.

**Supplementary Tables:**

Supplementary Table 1. Prior sleep and light exposure

Supplementary Table 2. MoCA domain scores at baseline and following FL and SL exposure

### Supplementary Figures

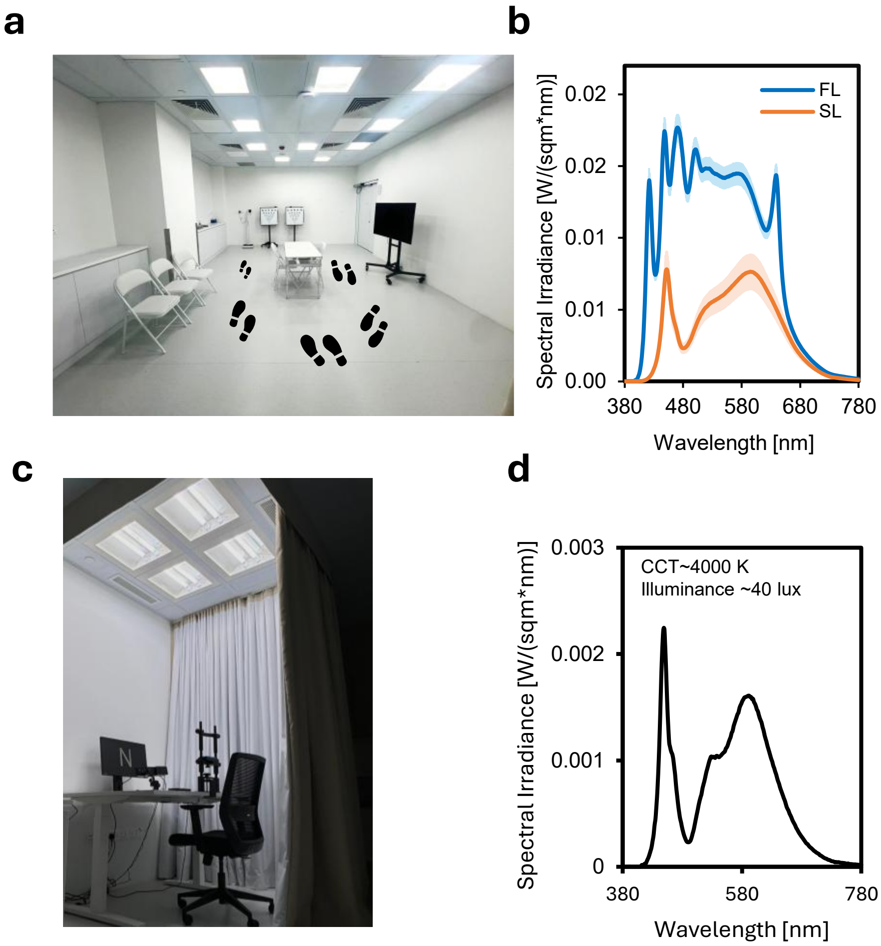


**Supplementary Fig. 1: Experimental setup and light source characterisation.**

**(a)** Light exposure room showing the ceiling-mounted full-spectrum light panels and a simulation of the walking circuit (footsteps) followed by participants during the exposure. **(b)** Spectral irradiance of the full-spectrum (FL, blue) and standard indoor (SL, orange) light conditions; shaded areas represent the measurement range across the room. **(c)** Cognitive testing station with the EyeLink 1000 Plus eye tracker and computer screen, enclosed by blackout curtains to isolate participants from the light exposure environment during task performance. **(d)** Spectral irradiance of the dim ambient light (~40 lux, CCT ~4000 K) present in the testing enclosure during cognitive assessments.

*CCT, Correlated color temperature; FL, full-spectrum light; SL, standard light.*

**a**

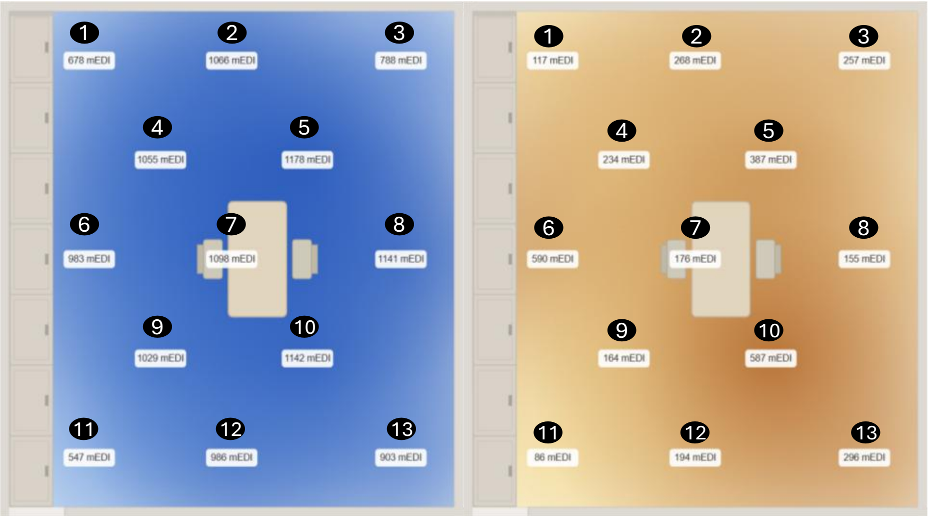

**b**

| | Standard Light (SL) · CCT ≈ 3955 K | | Full-Spectrum Light (FL) · CCT ≈ 6205 K | |
|---|---|---|---|---|
| | Lux (SL) | mEDI (SL) | Lux (FL) | mEDI (FL) |
| 1 | 190 | 117 | 705 | 678 |
| 2 | 436 | 268 | 1106 | 1066 |
| 3 | 417 | 257 | 813 | 788 |
| 4 | 383 | 234 | 1118 | 1055 |
| 5 | 628 | 387 | 1242 | 1178 |
| 6 | 956 | 590 | 1016 | 983 |
| 7 | 287 | 176 | 1132 | 1098 |
| 8 | 254 | 155 | 1172 | 1141 |
| 9 | 270 | 164 | 1078 | 1029 |
| 10 | 959 | 587 | 1189 | 1142 |
| 11 | 141 | 86 | 568 | 547 |
| 12 | 320 | 194 | 1023 | 986 |
| 13 | 492 | 296 | 940 | 903 |
| Median | 383 | 234 | 1078 | 1029 |
| IQR | 270-492 | 164-296 | 940-1132 | 903-1098 |

**Supplementary Fig. 2: Spatial distribution of melanopic equivalent daylight illuminance (mEDI) across the light exposure room.** (a) Heatmaps showing the spatial distribution of mEDI (lux) at 13 measurement positions across the room under the full-spectrum (FL, left, blue) and standard indoor (SL, right, orange) light conditions. Numbers indicate measurement positions; values represent mEDI in lux. (b) Photopic illuminance (lux) and mEDI at each of the 13 positions for both conditions, with median and interquartile range. All measurements were taken at 150 cm height facing upward using a calibrated spectroradiometer; mEDI was calculated per CIE S 026.

*CCT, Correlated color temperature; FL, full-spectrum light; SL, standard light.*

### Supplementary Tables

**Supplementary Table 1. Prior sleep and light exposure**

| Variable | Overall Median [IQR] |
|---|---|
| **Sleep — Median of 10 to 14 days** | |
| Mid-sleep time | 04:47 [04:16–05:34] |
| Time in bed (h) | 7.5 [7.1–7.8] |
| Total sleep time (h) | 6.5 [6.2–6.9] |
| Sleep efficiency (%) | 88.5 [86.0–91.2] |
| Onset latency (min) | 6.7 [4.7–8.2] |
| WASO (min) | 28.1 [18.8–38.7] |
| Number of Awakenings | 19.7 [12.6–24.4] |
| Inertia (min) | 12.6 [7.7–18.8] |
| **Light exposure — Median of 10 to 14 days** | |
| Outdoor time (min) | 96.1 [56.6–128.3] |
| Daytime lux | 204.3 [146.1–298.9] |
| Daytime mEDI (lux) | 169.0 [126.6–286.5] |
| Pre-sleep mEDI (lux) | 27.6 [16.5–44.9] |
| Sleep mEDI (lux) | 0.9 [0.2–6.1] |

*WASO: Wake after sleep onset*

**Supplementary Table 2. MoCA domain scores at baseline and following FL and SL exposure**

| Domain | Baseline[#] | SL | FL | Baseline vs SL | Baseline vs FL |
|---|---|---|---|---|---|
| Visuospatial / executive (/5) | 5.0 [4.0–5.0] | 4.0 [3.0–5.0] | 4.5 [4.0–5.0] | p = 0.005 ↓ | p = 0.15 |
| Naming (/3) | 3.0 [3.0–3.0] | 3.0 [3.0–3.0] | 3.0 [3.0–3.0] | ns | ns |
| Attention (/6) | 6.0 [6.0–6.0] | 6.0 [6.0–6.0] | 6.0 [6.0–6.0] | ns | ns |
| Language (/3) | 2.0 [2.0–3.0] | 2.0 [1.0–2.0] | 2.0 [2.0–3.0] | p = 0.02 ↓ | p = 0.61 |
| Abstraction (/2) | 2.0 [2.0–2.0] | 2.0 [2.0–2.0] | 2.0 [2.0–2.0] | ns | ns |
| Delayed recall (/5) | 5.0 [5.0–5.0] | 4.5 [4.0–5.0] | 5.0 [4.0–5.0] | p = 0.03 ↓ | p = 0.03 ↓ |
| Orientation (/6) | 6.0 [6.0–6.0] | 6.0 [6.0–6.0] | 6.0 [6.0–6.0] | ns | ns |
| Total score (/30) | 28.0 [27.0–29.0] | 27.0 [25.8–28.0] | 27.5 [26.0–28.2] | p = 0.02 ↓ | ns |

*Data are median [IQR], N = 24. Wilcoxon signed-rank test (two-tailed). ↓ = significantly lower than baseline; ns = not significantly different than baseline. FL: full-spectrum light; SL: standard light.*
*[#] Baseline scores were obtained at Visit 1 using the Singapore culturally adapted MoCA version, which differs from the English versions (V8.2/V8.3) administered at Visit 2. Comparisons between baseline and Visit 2 conditions are therefore exploratory and should be interpreted with caution.*
